# First record of *Argulus japonicus* infestation on *Cyprinus carpio* in Hungary, and the first description of *Argulus japonicus* subsp. *europaeus* subsp. nov. Keve, 2025

**DOI:** 10.1101/2025.04.09.647856

**Authors:** Gergő Keve, Adrienn Gréta Tóth, Máté Katics, Ferenc Baska, Edit Eszterbauer, Sándor Hornok, Tibor Németh, Norbert Solymosi

## Abstract

Species belonging to the genus *Argulus* are globally distributed fish parasites. Their veterinary significance primarily lies in their disruptive presence and their role as mechanical vectors. Although *Argulus japonicus* Thiele, 1900 is a widely distributed representative of this genus that feeds on freshwater fish, only *Argulus foliaceus* (Linnaeus, 1758) had previously been reported in Hungary. The aim of this study was to investigate the fish louse fauna in a local common carp (*Cyprinus carpio* Linnaeus, 1758) population. To the best of our knowledge, this is the first study to report the occurrence of *A. japonicus* in Hungary. Our detailed molecular analyses, including the complete mitochondrial genome, revealed for the first time that the *A. japonicus* specimens found in Hungary differ significantly from their Far Eastern counterparts. Furthermore, cytochrome c oxidase subunit I (*cox1*) sequence analysis — a region known to be stable within the species — showed that while our sequences were nearly identical to those of other European specimens, they differed markedly from the available Asian isolates. The phylogenetic analysis also confirmed this divergence. The European *A. japonicus* sequences form a clearly distinct sister group to the Asian lineages. In light of these findings, and based on thorough morphological examinations, the authors propose that the specimens found in Hungary represent a new subspecies, *Argulus japonicus* subsp. *europaeus* subsp. nov. Keve, 2025.

## Introduction

Fish lice belonging to the Argulidae family are a well-known group of crustacean fish parasites. Only three freshwater species of the family have been confirmed in Europe: *Argulus coregoni* (Thorell, 1864), *Argulus foliaceus* (Linnaeus, 1758) and *Argulus japonicus* Thiele, 1900 (Fryer, 1982).^1^ While fish lice occasionally parasitize amphibians or invertebrates, their primary hosts are fish,^2^ including ones of high economic importance, such as cyprinids (*Cyprinus carpio, Chondrostoma* sp., *Squalius cephalus*). The presence of fish lice is disadvantageous at fish farms.^3,4^ This is due to their parasitic nature that causes constant stress to the fish and their role as mechanical vectors for the transmission of various pathogenic agents, such as the spring viremia of carp virus (SVCV).^5^ At the same time, knowledge on the distribution range of *Argulus* species is not complete. The morphological differences between the adults of various *Argulus* species are not always pronounced. Moreover, they have multiple developmental stages, each with somewhat different morphological characteristics.^6^ On the other hand, with the rise of molecular identification methods, e.g. species identification based on specific gene sequences, like *cox*1 new doors have opened for taxonomists. Today, the identification of new species should contain morphological and also molecular approaches. To date, the scientific literature has documented only one species of fish lice, *A. foliaceus* being present in Hungary.^7–9^ The primary objective of the present study was to gain insight into the presence and range of pathogens associated with *Argulus* infestation at a fish farm in Hungary. Thus, next-generation shotgun sequencing (NGS) based metagenomic analysis was performed using the total DNA extracted from fish lice. By such NGS-based metagenomic studies the vast majority of the generated sequences derive from the host genome. Thus, the approach is applicable to gain insights into the taxonomic properties of the host, fish lice and perform its classification. The genomic results from the initial metagenomic sampling were complemented with additional samples of fish lice from the same fish farm in order to evaluate its taxonomy with the classical morphological analysis as well.

## Materials and Methods

### Sampling and morphology

The first sample collection was performed on 05/04/2024, and the second set of samples was gathered on 07/10/2024. The samples were stored at -20^°^C.

For morphological identification, *Argulus* specimens were put into saline and were observed with stereo (Leica M205C) and with light microscopes (Leica DM2000) (Leica Microsystems, Wetzlar, Germany). The morphological identification of *A. japonicus* was based on morphological keys and on relevant articles^1,10–14^. Photography for Figure 1 and Figure 2 were made with a stereo microscope and compatible camera Leica DMC4500, and were compiled and measured with LasX (Leica application suite X) program (Leica Microsystems, Wetzlar, Germany). Photography for Figure 3, Figure 4, Figure 5, Figure 6 were made with light microscope and compatible camera Leica FLEXACAM C1 (Leica Microsystems, Wetzlar, Germany) and were compiled with CombineZP program^15^. For measurements, both microscopes were calibrated with the same objective micrometer. Scales were added or adjusted manually.

**Figure 1.**
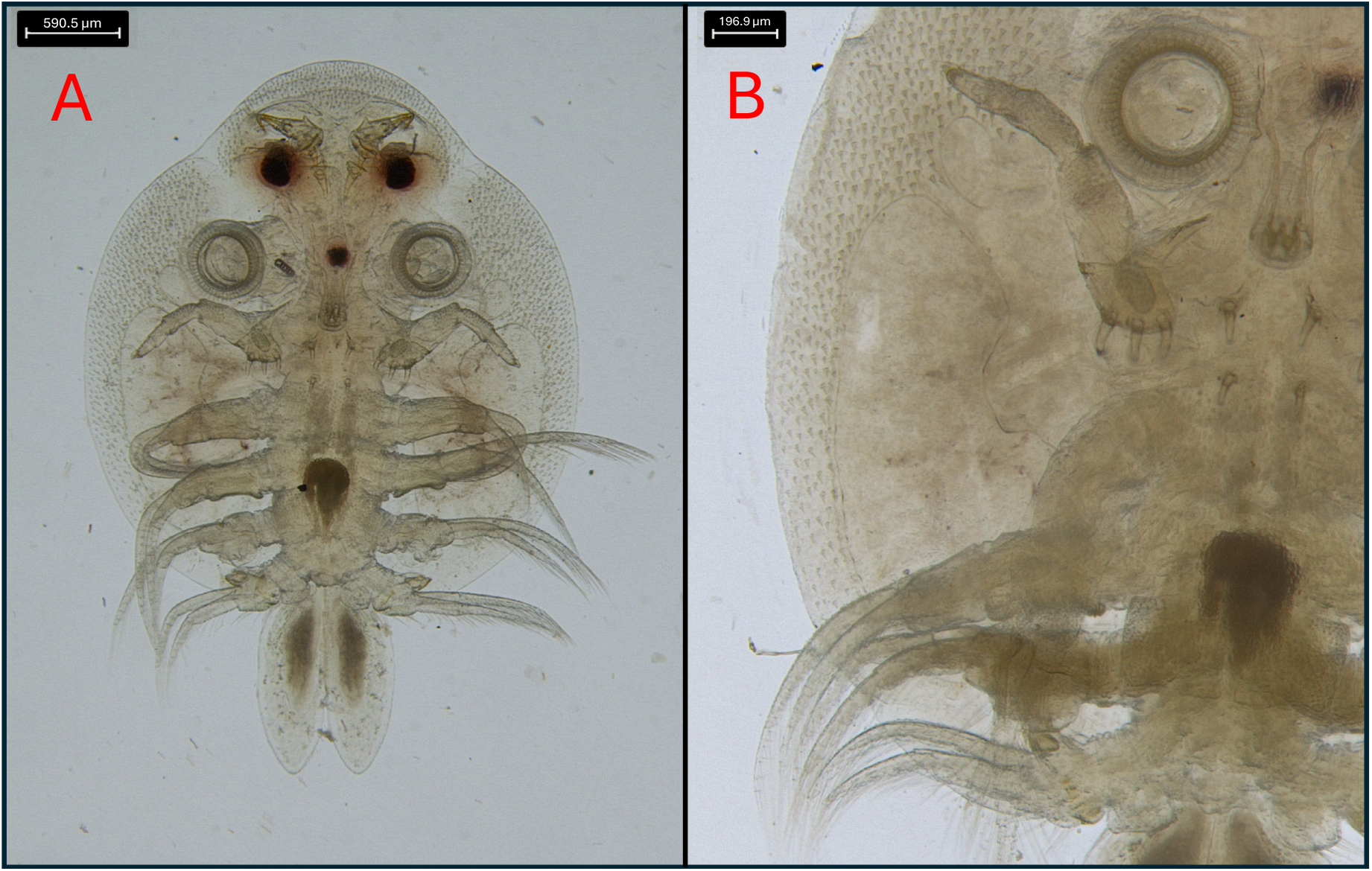
A) Ventral aspect of an *A. japonicus* subsp. *europaeus* subsp. nov. male; B) The respiratory areas of the same specimen.

**Figure 2.**
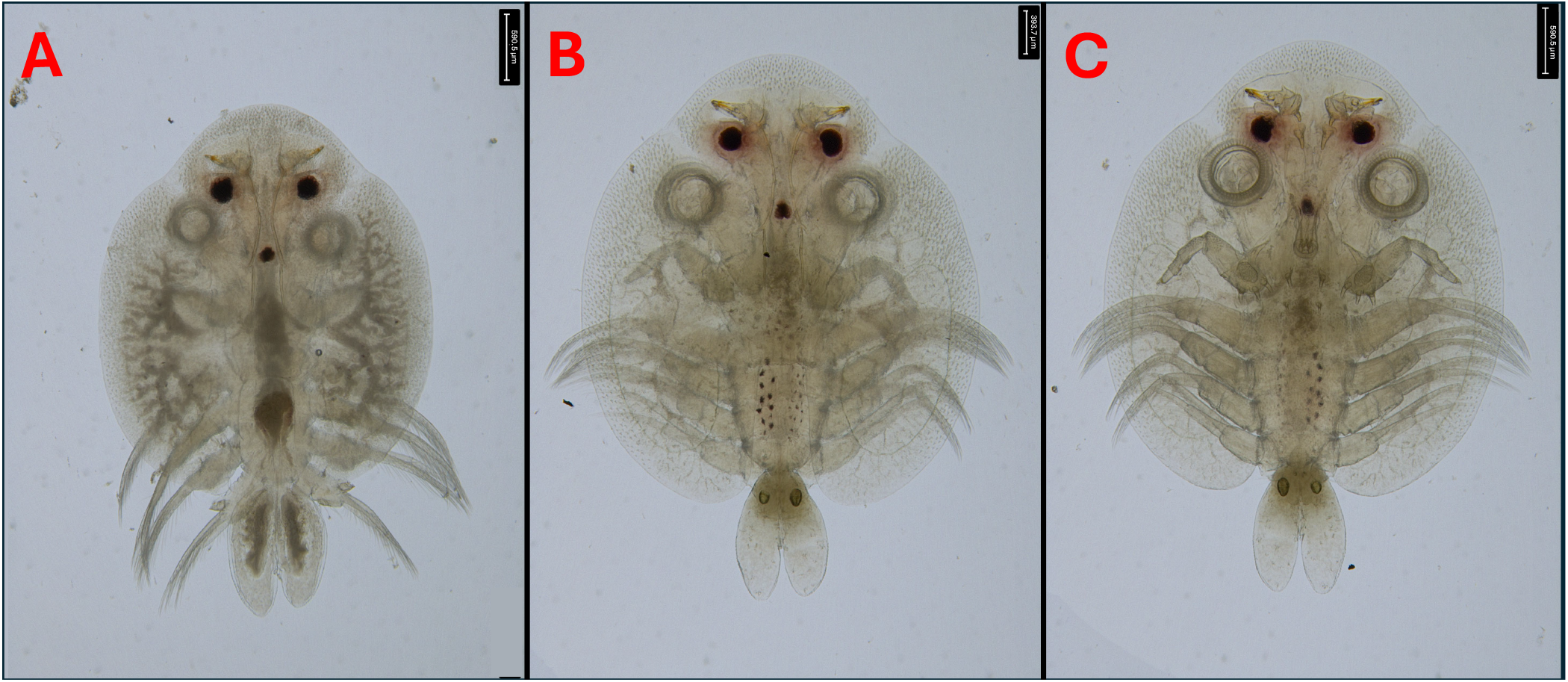
A) Dorsal aspect of a male (this specimen is different from the one on Figure 1); B) Dorsal aspect of a female; C) Ventral aspect of a female *A. japonicus* subsp. *europaeus* subsp. nov.

**Figure 3.**
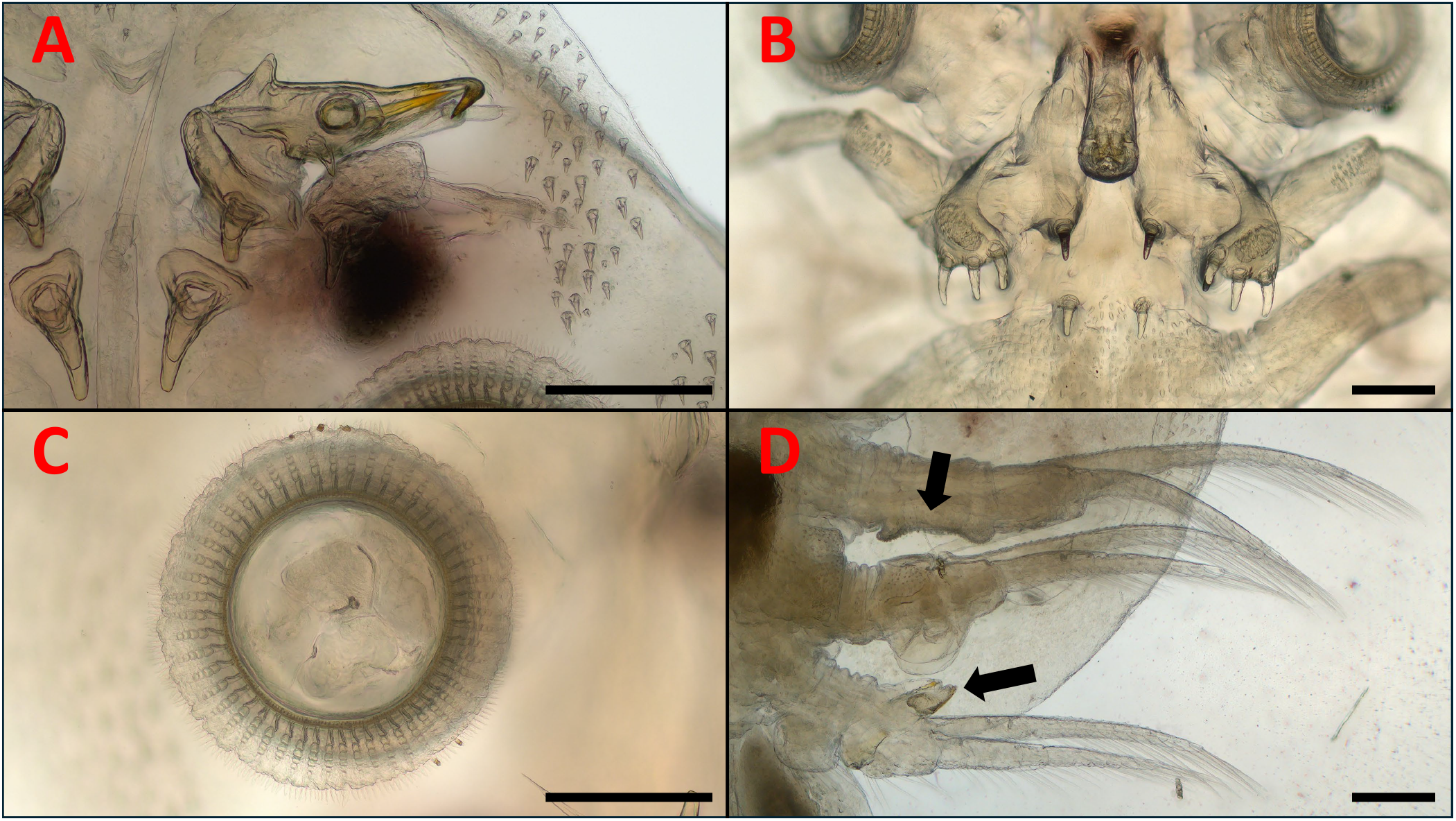
The relevant morphological structures of an *A. japonicus* subsp. *europaeus* subsp. nov. male specimen. A) Antennae and stylet; B) Secondary maxillae and mouth tube (50x magnitude); C) Sucker and its supporting rods; D) The 2nd, 3rd and 4th legs (arrows: clasping apparatus on legs 2 and 4, characteristic to *A. japonicus*. Scales: 250 µm

**Figure 4.**
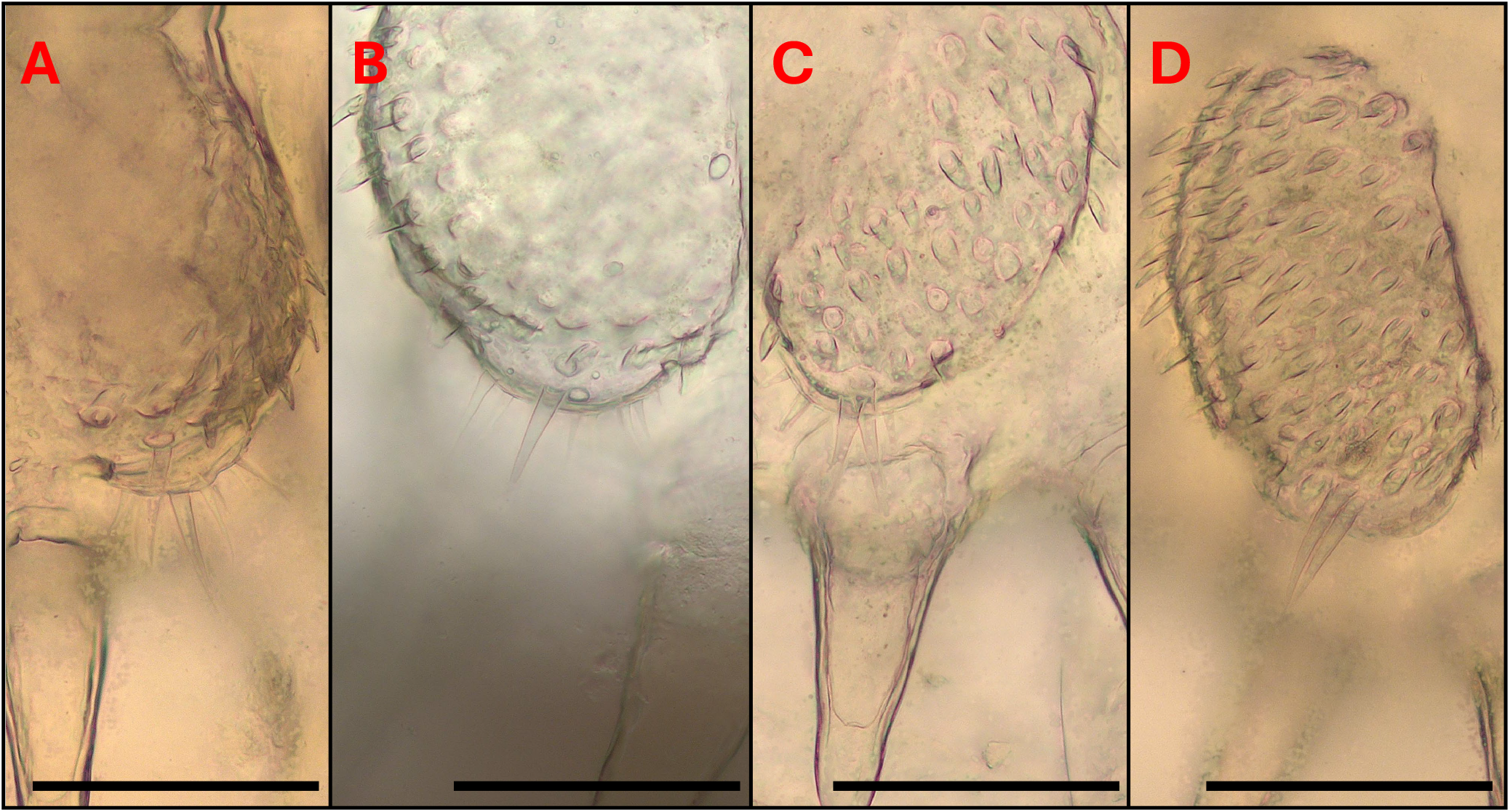
Setae on the margin of the secondary maxillae of females (A, B) and males (C, D) scales: 100 µm

**Figure 5.**
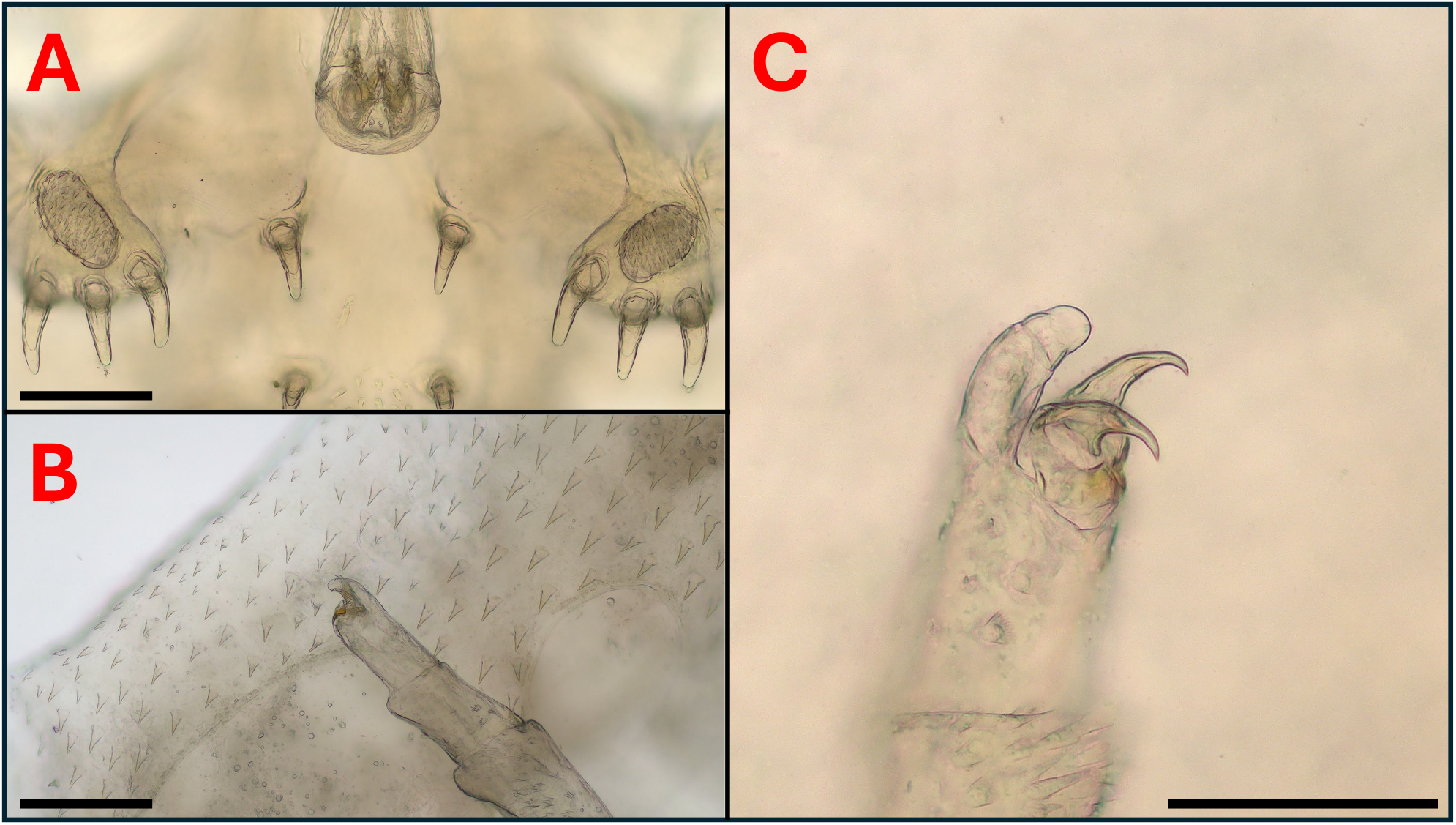
A) Labium and secondary maxillae of a male; B) Final segment of the secondary maxilla of a female; C) Final segment of the secondary maxilla of a male. Scales: 100 µm

**Figure 6.**
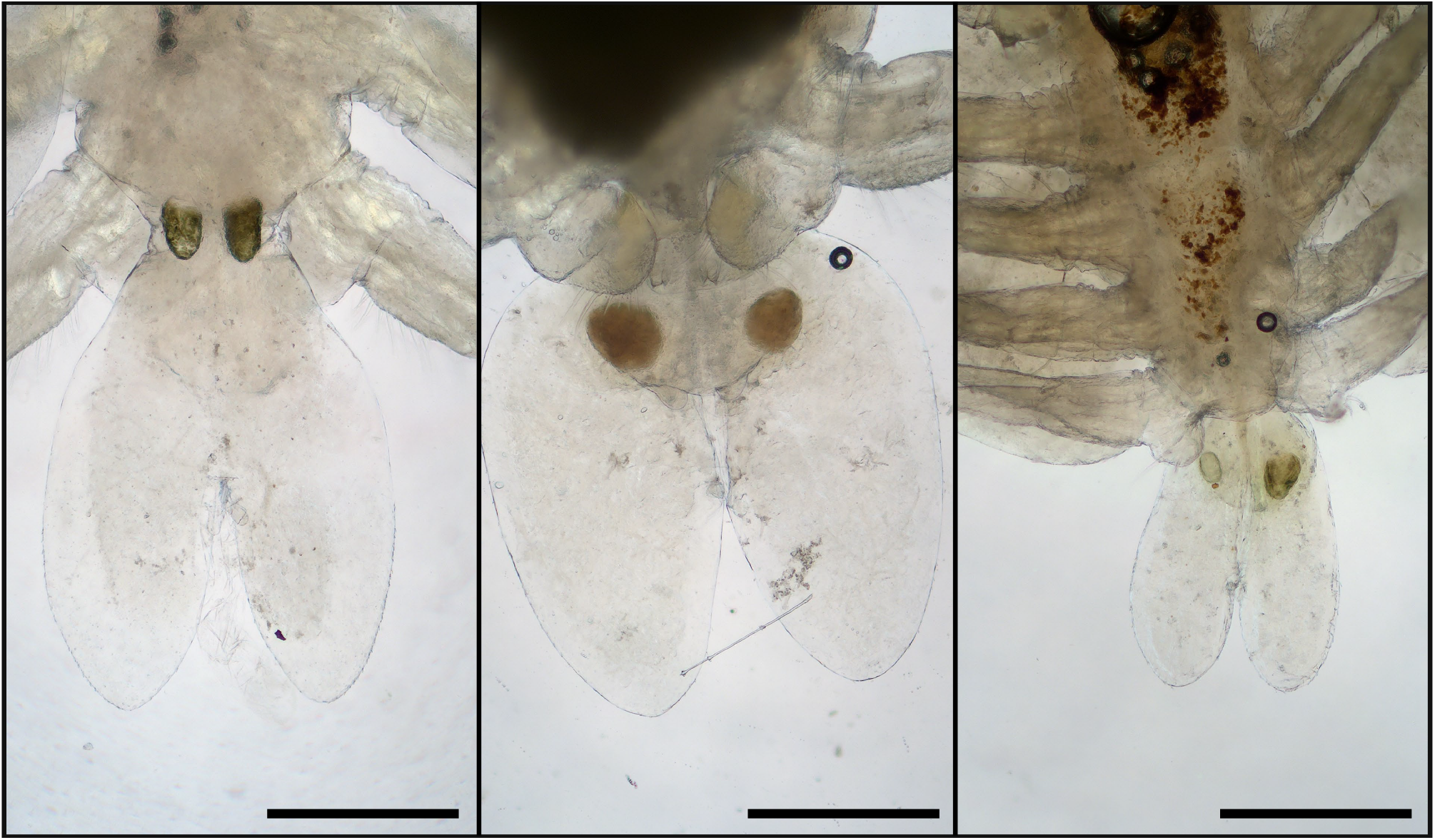
Abdominal lobes of three different female specimens. Scales: 500 µm

### Sequencing

DNA purification from the sample was performed in triplicates, and the resulting total DNA extracts were pooled together. DNA extraction for Illumina and Nanopore sequencing were carried out using ZymoBIOMICS DNA/RNA miniprep kits (R2002, Zymo Research, Irvine, USA). For efficient sample lysis, bead homogenization was performed using a Vortex-Genie 2 with a bead size of 0.1 mm and a homogenization time of 15 min at maximum speed. After that, the Zymo Research kit’s DNA purification protocol was followed. Total DNA qualities were assessed with an Agilent 2200 TapeStation instrument (Agilent Technologies, Santa Clara, USA), and DNA quantities were measured using a Qubit Flex Fluorometer (Thermo Fisher Scientific, Waltham, MA, USA). We closely followed all manufacturer recommendations when preparing sequencing libraries for Illumina sequencing platform (Illumina Inc., San Diego, USA). Pooled total DNA samples were constructed using the NEBNext Ultra II Library Prep Kit (NEB, Ipswich, MA, USA). Paired-end shotgun metagenome sequencing was performed on a NextSeq 550 (Illumina, San Diego, CA, USA) sequencer using the NextSeq High Output Kit v2 sequencing reagent kit. Primary data analysis (i.e., base-calling) was performed using “bcl2fastq” software (version 2.17.1.14, Illumina).

For sanger sequencing, the specimens (two males, including the one in Figure 1, and two females) were disinfected on their surface with sequential washing for 15 s in detergent, tap water, and distilled water. For the DNA extraction, legs and pieces of the carapace were cut off and used. DNA was extracted with the QIAamp DNA Mini Kit (QIAGEN, Hilden, Germany) according to the manufacturer’s instructions, including an overnight digestion in tissue lysis buffer and Proteinase-K at 56 °C. Extraction controls (tissue lysis buffer) were also processed with the *Argulus* samples to monitor cross-contamination.

The *cox*1 gene was chosen as the first target for molecular analysis. The PCR was modified from Folmer et al.^16^ and amplifies an approx. 710-bp-long fragment of the gene. The primer LCO1490 (5’-GGT CAA CAA ATC ATA AAG ATA TTG G-3’) were used in a reaction volume of 25 µl, containing 1 U (stock 5 U/ µl) HotStarTaq Plus DNA Polymerase, 2.5 µl 10× CoralLoad Reaction buffer (including 15 mM MgCl_2_), 0.5 µl PCR nucleotide Mix (stock 10 mM), 0.5 µl of each primer (stock 50 µM), 15.8 µl ddH_2_O and 5 µl template DNA. For amplification, an initial denaturation step at 95°C for 5 min was followed by 40 cycles of denaturation at 94°C for 40 s, annealing at 48°C for 1 min and extension at 72°C for 1 min. Final extension was performed at 72°C for 10 min.

In all PCRs non-template reaction mixture served as negative control. Extraction controls and negative controls remained PCR negative in all tests. Purification and Sanger sequencing of the PCR products were done by Biomi Ltd. (Gödöllő, Hungary).

### Bioinformatic analysis

For raw Illumina sequenced data, the quality-based filtering and trimming of the raw short reads were performed by TrimGalore (v.0.6.6, https://github.com/FelixKrueger/TrimGalore), setting 20 as the quality threshold. Only reads longer than 50 bp were retained. Using default settings, the cleaned reads were assembled to contigs by MEGAHIT (v1.2.9)^17^.

The basecalling was performed using dorado (https://github.com/nanoporetech/dorado, v0.9.0) with model dna_r10.4.1_e8.2_400bps_sup@v5.0.0, based on the POD5 files generated by the ONT Mk1C sequencer. The raw reads were adapter-trimmed and quality-based filtered by Porechop (v0.2.4, https://github.com/rrwick/Porechop) and Nanofilt (v2.6.0, minimal Q=7, length=50)^18^, respectively.

The assembled contigs and the ONT long reads were taxon classified by Kraken2 (v2.1.4)^19^ using the NCBI Core NT database (created: 12/28/2024). The *Argulus* hits were analysed by BLAST^20^ on NCBI Core NT database (accessed: 01/03/2025). The mitogenomes were annotated and visualized by MitoZ.^21^ The identity of detected mitogenome features was analyzed in the R-environment^22^ by the Needleman-Wunsch global alignment algorithm of package pwalign.^23^ Phylogenetic analysis was performed based on the COX1 overlapping region. The gene-tree was constructed^24^ based on multiple sequence alignment by MAFFT (v7.490).^25^ The best substitution model was selected by functions of phangorn (v2.11.1) package^26^ based on the Bayesian information criterion. The generated neighbor-joining tree was optimized by the maximum likelihood method. Bootstrap values were produced by 100 iterations. All data processing and plotting were done in the R-environment.^22^ Quality control and trimming of sequences were performed done with the BioEdit program (v7.7.1.0), then alignment with GenBank sequences by the nucleotide BLASTN program The analyses of assembled sequences were performed with BLASTN via GenBank (https://blast.ncbi.nlm.nih.gov).

## Results

According to our morphological analyses, the fish lice we found belong to the *A. japonicus* species: The coxae with the clasping apparatus of the second, third, and fourth legs of the males have the same appearance as described in the work of Fryer.^10^ (Figure 1/A, Figure 3/D). Other works, such as those of Rushton-Mellor and Boxshall, and Shoes et al.^1,6^ also underline, that the most reliable morphological attributes based on which *A. japonicus* and *A. foliaceus* can be distinguished are the different accessory copulatory structures on legs 2, 3, and 4 of the males.

In the case of both males and females, the abdominal lobes are acutely rounded, as described in the work of Fryer^10^, although, according to our observations, their general shapes in females were variable (Figure 6). The larger respiratory areas are reniform and posterior to the smaller respiratory areas (Figure 1/B). On the antennae, distinct, sharp terminal, anterior, medial, posterior, and post-antennal spines are visibe (Figure 3/A).

Basal plates are present on the second maxillae. The coxal spines of the secondary maxillae are long and finger-like (Figure 3/B). Scales are present on the labium, although rather subtle (Figure 5/A).

The basal sclerites of the supporting rods of the suckers are elongated (number of supporting rods: 52-53 in males and females, with 5-7 sclerites in each rod. (Figure 3/C). This is more or less in line with the observations of Nagasawa^13^ (50-52 rods in a female) and those of Wadeh et al.^12^ (45-53 rods). The hooks on the final segments of the second maxillae of both females and males are similar to the figures in the work of Nagasawa^13^ (Figure 5/B,C).

Relevant differences were observed in some specimens, compared to the works of Nagasawa^13^ and Rushton-Mellor^27^: the post antennal spine on our specimens have widened basis in contrast to the specimens described in the aforementioned works Figure 3/A. On the margin of the basal plate of the secondary maxilla, there are two long setae, with several medium-long setae (instead of just two long setae) (Figure 4) A/B. On the other hand, among males, some specimens had two setae, while others possessed multiple, just as the females (Figure 4C/D). However, in the work of Bauer^11^, these medium-length setae are also reported in the case of *A. japonicus*.

The abdominal incision of the female is less than half of the abdominal length in contrast to the description of Fryer^10^ (Figure 6. The total body length of females was in the range of *A. japonicus* 3.12-6.27 mm (mean = 4.67, n = 12), while the length of males was between 3.81-5.49 mm (mean = 4.55, n = 9). Even though these measurements differ from the sizes reported by Wadeh et al.^12^ (Figure 2/A,B,C), a comparison should not be made, since these crustaceans were potentially in different life stages.

We assembled an MT-derived contig from the Illumina sequencing, while a long read from the ONT sequencing was found to be MT-derived.

The length of the assembled MT-sequence of Arg_hun1 sample reached 15004 bp. Detailed annotation of this sequence can be found in Table 1 and Figure 8. Since only *A. foliaceus* has been scientifically proven to be present in Hungary, the first comparison aimed at this species as the reference sequence for our MT sequence. However, no complete *A. foliaceus* sequences are available in the NCBI NT database, only shorter fragments of it. Consequently, the basis of the comparison relied on common, incomplete MT-sequences that overlapped in multiple *Argulus* species, encompassing the 10810-11442 region of our assembled MT-sequence. This segment showed the highest sequential similarity to *A. japonicus* (OL841710.1) *cox*1 (identity: 565/565 (100%), gaps: 0/565 (0%)), escorted by the following hits: *A. mongolianus* (OL841700.1, identity: 467/574 (81%), gaps: 0/574 (0%)), *A. coregoni* (OL841703.1, identity: 461/565 (82%), gaps: 0/565 (0%)), *A. americanus* (NC_005935.1, identity: 494/633 (78%), gaps: 0/633 (0%)), *A. siamensis* (KF713308.1, identity: 486/633 (77%), gaps: 0/633 (0%)), *A. foliaceus* (KF723419.1, identity: 478/633 (76%), gaps: 0/633 (0%)), and *A. yucatanus* (ON715940.1, identity: 411/528 (78%), gaps: 0/528 (0%)). The complete MT-sequence was associated with higher sequential identity to *A. japonicus* (LC588400.1: coverage: 98%, identity: 84.82%; NC_088557.1: coverage: 98%, identity: 84.68%) than with *A. americanus* (NC_005935.1: coverage: 90%, identity: 76.58%).

**Table 1.**
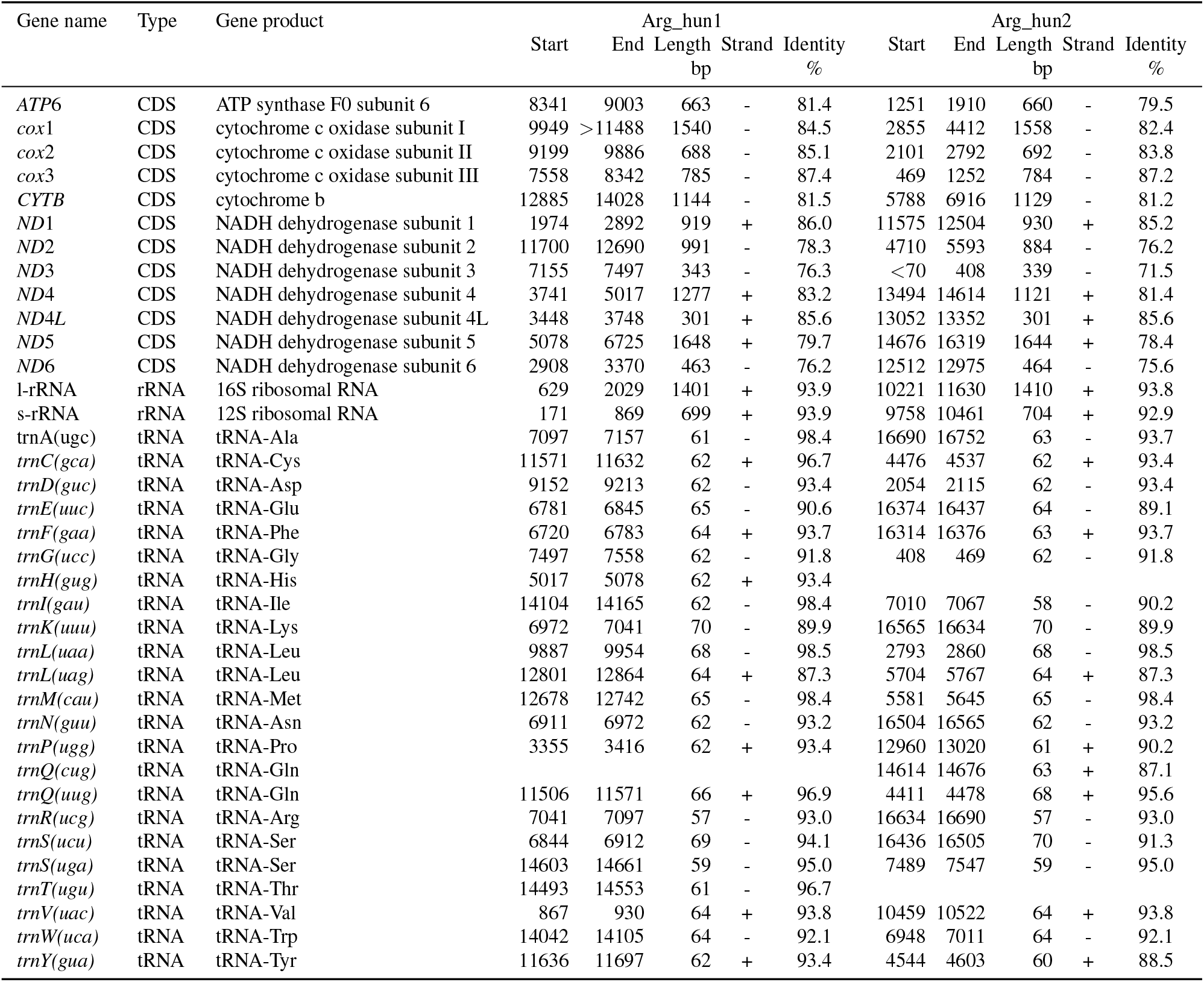
Annotation of mitogenome the two sequenced samples. The sequential identity of the detected 37 features with the reference genome (LC588400.1) features was calculated by Needleman-Wunsch global alignment.

**Figure 7.**
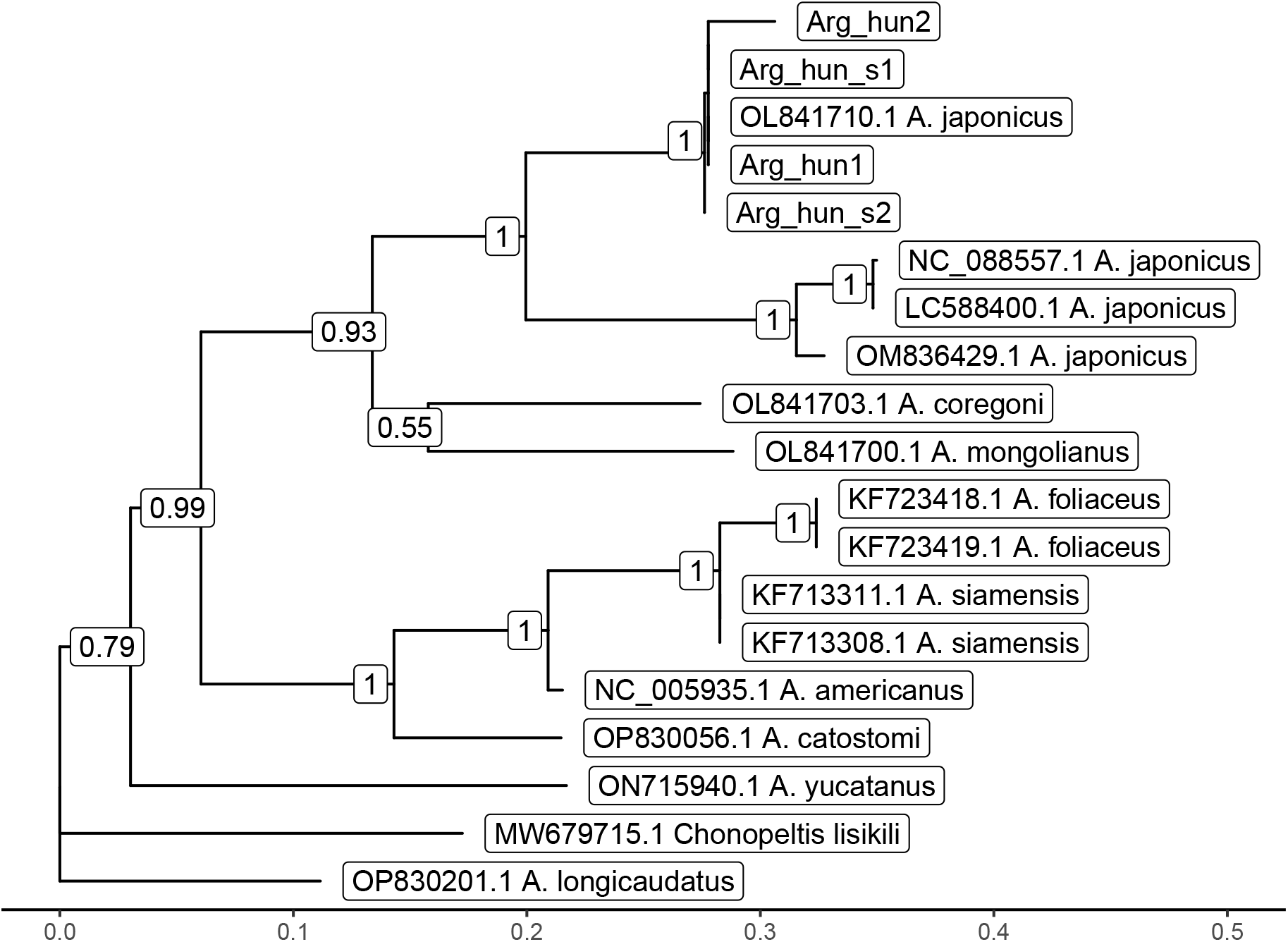
Gene tree based on the overlapping segment of *cox*1. Outgroup: *Chonopeltis lisikili* (MW679715.1). Numbers at branches indicate bootstrap support levels (100 replicates).

**Figure 8.**
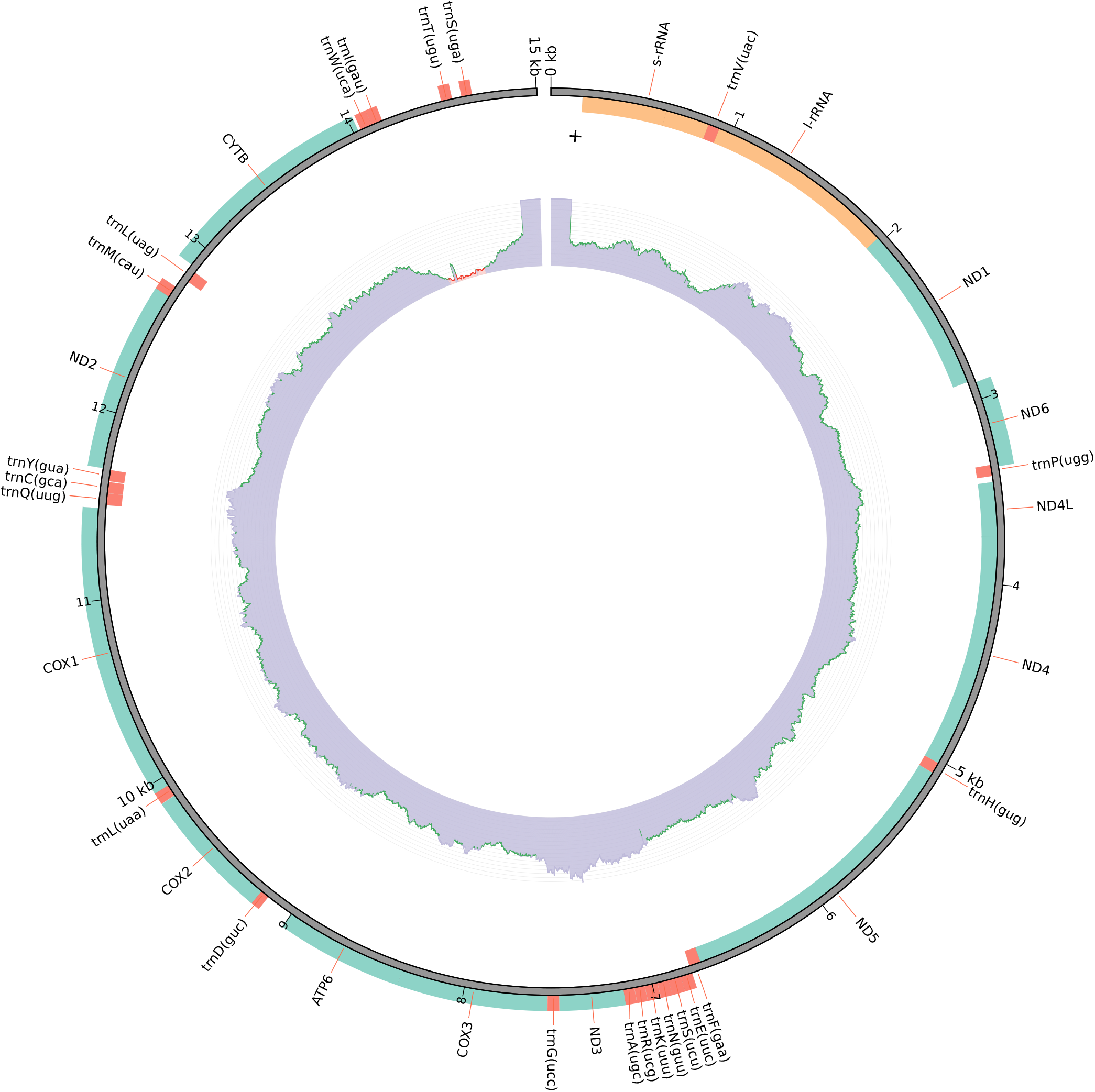
Mitogenome visualization of Arg_hun1 sample. The outer circle shows the genomic features (n=36) identified, and the inner one traces the depth of mitogenome coverage.

The 619 bp length *cox*1 sequences of one male and two females received after Sanger sequencing were 100% identical to European *A. japonicus* sequences: OL841708.1 - OL841710.1 (565/565 (100%), gaps: 0/565 (0%)) as well. These sequences showed a low sequential similarity 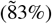 to Japanese and Chinese *A. japonicus* sequences (LC588400.1 and NC_005935.1; 513 and 512/618 bp identity, respectively, with 0 gaps). One male specimen differed from the rest by only one nucleotide (618/619 bp (99.84%)

The length of the MT-sequence of Arg_hun2 sample reached 16797 bp. Detailed annotation of this sequence can be found in Table 1 and Figure 9. The same segment as above showed the highest sequential similarity to *A. japonicus* (OL841710.1) *cox*1 (identity: 545/569 (96%), gaps: 12/569 (2%)), escorted by the following hits: *A. mongolianus* (OL841700.1, identity: 456/578 (79%), gaps: 14/578 (2%)), *A. coregoni* (OL841703.1, identity: 447/568 (79%), gaps: 5/568 (1%)), *A. americanus* (NC_005935.1, identity: 484/639 (76%), gaps: 15/639 (2%)), *A. longicaudatus* (OP830201.1, identity: 480/634 (76%), gaps: 13/634 (2%)), *A. catostomi* (OP830056.1, identity: 482/637 (76%), gaps: 13/637 (2%)), *A. siamensis* (KF713311.1, identity: 476/638 (75%), gaps: 13/638 (2%)), *A. foliaceus* (KF723419.1, identity: 468/638 (73%), gaps: 13/633 (2%)), and *A. yucatanus* (ON715940.1, identity: 402/533 (75%), gaps: 10/533 (1%)). The complete MT-sequence was associated with higher sequential identity to *A. japonicus* (LC588400.1: coverage: 87%, identity: 83.71%; NC_088557.1: coverage: 87%, identity: 83.49%) than with *A. americanus* (NC_005935.1: coverage: 79%, identity: 75.26%).

**Figure 9.**
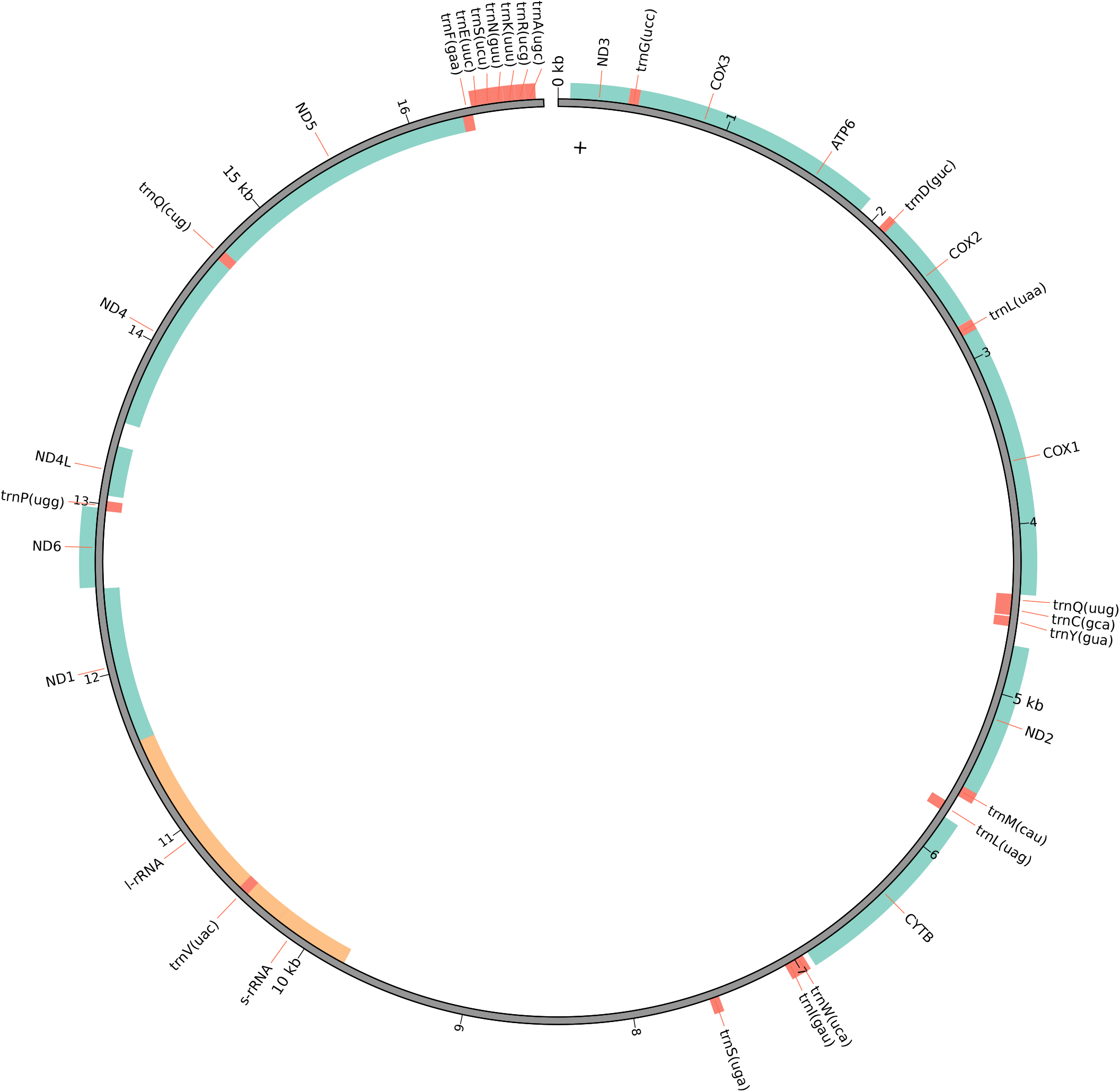
Mitogenome visualization of Arg_hun2 sample. The circle shows the genomic features identified.

The BLAST alignment of the Arg_hun1 MT-DNA sequence on Arg_hun2 gives 100% coverage and 97.83% sequential identity. Figure 7 shows the gene-tree based on the above-mentioned overlapping *cox*1 sequences with the best substitution model, TVM. On this tree, it is visible that *A. japonicus* samples from Asia (China, Japan, and India) form a highly distinct sister group compared to European samples (Hungary and England), with strong statistical support (1).

Based on the morphological, and genetic differences compared to the (Asian) *A. japonicus* samples, it is assertable that the Hungarian *A. japonicus*, as well as some other European specimens reported before belong to a new subspecies: *Argulus japonicus* subsp. *europaeus* subsp. nov. Keve, 2025. The holotype (male specimen, collected by Adrienn Gréta Tóth and Norbert Solymosi on 07/10/2024, Hungary) is deposited in the Department of Parasitology and Zoology, University of Veterinary Medicine, Budapest (accession number: UNIVET-PAR-KG001.) ZooBank registration: To comply with the regulations set out in article 8.5 of the amended 2012 version of the International Code of Zoological Nomenclature (ICZN)^28^, details of the new subspecies have been submitted to ZooBank. The Life Science Identifier (LSID) of the article is LSIDurn:lsid:zoobank.org:pub:27F531B8-87D0-4CE3-A98C-6D526E6D7795. The LSID for the new name *Argulus japonicus* subsp. *europaeus* Keve, 2025 is LSIDurn:lsid:zoobank.org:act:4D58B113-04B5-472A-AA8E-A015F5AA8449. The sequences obtained in the current study were deposited in the GenBank database and are available under the following accession numbers.

## Discussion

*Argulus japonicus* was originally described in China. Besides its Asian distribution including Bangladesh, China, India, Japan, Syria, and Turkey,^12,29–33^ its global spread is also reported. Its appearance is recorded in Africa, Australia, Europe, North America, and South America.^12,29,34–37^ However, its precise distribution range is less studied and several countries lack information on the presence of the parasite. In Europe, *A. japonicus* was described in Bosnia and Herzegovina, Croatia, Montenegro, France, Germany, Greece, Italy, the Netherlands, Norway, Poland, the United Kingdom, Serbia, Slovakia, and Spain.^1,38–43^ Although many of these reports are related to fish trading and imported ornamental fish, several findings are derived from fish farms or wild waters.

To the best of our knowledge, this is the first report confirming the presence of *A. japonicus* in Hungary and the first study to provide the complete mitochondrial genome of a European *A. japonicus* specimen. However, this 15,004 bp sequence exhibits low similarity (85%) to previously reported Japanese (LC588400: identity 9627/11313 [85%], gaps 99/11313 [0%]) and Chinese (NC_088557: identity 9620/11324 [85%], gaps 117/11324 [1%]) sequences. Also, based on their morphology and *cox1* sequences, these fish-lice we found are identical to other *A. japonicus* found in Europe, and differ from *A. foliaceus*, a species that is known to be present in Hungary (Figure 7). This is despite the fact, that the *cox*1 gene is reported to be relatively stable in the case of the species *A. japonicus* from Asia and Africa (variations are between 0.0-1.9%)^29^, and also suitable for species differentiation^44^.

Notably, our phylogenetic analysis based on the *cox*1 gene reveals that *A. japonicus* samples from Asia (China, Japan, and India) form a highly distinct sister group compared to European samples (Hungary and England), with strong statistical support (1) (Figure 7). This finding raises an important question: Are the *A. japonicus* specimens reported in various European countries truly the same species as the *A. japonicus* commonly found parasitizing freshwater fish in Asia? In this study, we highlight the morphological similarities between these lineages while also addressing their genetic diversity. Based on our findings, we strongly suspect that *A. japonicus* represents a species complex with distinct lineages in Europe and Asia, rather than a single species. To determine the extent of this ectoparasite’s presence, comprehensive studies would be required to evaluate its distribution and frequency across Hungary. Additionally, given that no data is available on the simultaneous occurrence of the fish lice species, further sample collection and co-occurrence studies of *A. foliaceus* and *A. japonicus* could enhance our understanding of the infestation risk and possible interbreeding. Nevertheless, based on morphological and molecular differences, we recognize the fish lice discussed in this study as a new subspecies of *A. japonicus*, namely *A. japonicus* subsp. *europaeus* subsp. nov. Keve 2025.

## Declarations

### Ethics approval and consent to participate

Not applicable.

### Consent for publication

Not applicable.

### Availability of data and material Competing interests

The authors declare that they have no competing interests.

### Funding

The study was supported by the strategic research fund of the University of Veterinary Medicine Budapest (Grant No. SRF-001), and the European Union project RRF-2.3.1-21-2022-00004 within the framework of the MILAB Artificial Intelligence National Laboratory. Financial support was provided by the Office for Supported Research Groups, Hungarian Research Network (HUN-REN), Hungary (Project No 1500107).

### Author contributions statement

AGT, GK and NS conceived the concept of the study. AGT, GK, MK, and NS collected the samples. TN, SH provided the sequencing instrumentation and environment. SH assisted in species description and performed quality control of the Sanger sequencing results. AGT and GK did the laboratory work and sequencing. GK performed the morphological identification of the *Argulus* samples. GK performed the photography of the samples, FB and EE validated the morphology-related results, NS participated in the bioinformatic analysis. NS takes responsibility for the genomic data’s integrity and the data analysis’s accuracy. AGT, GK and NS participated in the drafting of the manuscript. AGT, GK, FB, EE, SH and NS carried out the critical revision of the manuscript for important intellectual content. All authors read and approved the final manuscript.

## Acknowledgements

The authors would like to thank László Békési, Géza Szita, and László Czikk for supporting their works on fish health.

## Notes

### Competing Interest Statement

The authors have declared no competing interest.

